# Scaling Quantum Optimisation Beyond Hardware Limits for Real-World Scientific Workloads: Genome Assembly on Current Quantum Hardware

**DOI:** 10.64898/2026.09.04.749434

**Authors:** Namasi G Sankar, Georgios Miliotis, Simon Caton

## Abstract

Genome assembly is important in infectious disease surveillance, antimicrobial resistance monitoring, and cancer genomics. The task of reconstructing full genomic sequences from fragmented reads, can be framed as a large scale combinatorial optimisation problem. Recent advances in quantum computing have introduced new optimisation algorithms with potential advantages for navigating complex combinatorial search spaces. However, practical deployment is limited by noisy intermediate-scale quantum (NISQ) hardware, including restricted qubit counts, limited connectivity, and high error rates. In this research, we employ the Hamiltonian Auto Decomposition Optimisation Framework (HADOF), an algorithm agnostic framework that enables scalable quantum optimisation through federated solving across small subproblems. HADOF enabled the quantum-assisted genome assembly of a 7.1 Million base pairs *Pseudomonas aeruginosa* genome, to our knowledge, representing the largest genome assembly graph studied on real quantum hardware to date. The results achieved a 99.348% genome fraction and 1.0 duplication ratio, demonstrating that biologically plausible genome reconstructions can be obtained despite current hardware limitations.

## 1 Introduction

Genome assembly is a fundamental task in genomics, aiming to reconstruct an organism’s complete DNA sequence from sequencing reads [44]. Sequencing reads are DNA fragments reported by a sequencing instrument and record the order of nucleotide bases (A, C, G and T) across part of a genome, but a single read typically covers only a small fraction of the full genome [51]. Read length depends on the sequencing platform. Second generation Illumina sequencing usually produces short reads in 50–300 base pair (bp) fragments. Conversely, third generation sequencing platforms such as PacBio (Pacific Biosciences) and Oxford Nanopore Technologies (ONT) produce longer 1-100+ kilo bp reads [45]. Genome sizes, however, vary enormously across the tree of life, ranging from under 10,000 bp in small viruses to over 100 billion bp (Gbp) in certain plants and amoebas. Sequencing reads capture fragments of the original genome and must be assembled to reconstruct the full sequence. The reconstruction can be done either by reference-guided assembly, which requires a suitable reference genome and prior knowledge that the sample is sufficiently related to, or by de novo assembly, which reconstructs the genome directly from read-to-read relationships without relying on an external reference [51]. This research focuses on de novo genome assembly using ONT long reads.

Accurate and timely genomic reconstruction has important applications in both clinical medicine and public health. In infectious disease surveillance, reconstructing complete pathogen genomes enables accurate species and strain identification, allowing researchers and clinicians to distinguish between closely related variants that may differ in virulence or antimicrobial resistance [11]. Comparing assembled genomes from different patients also makes it possible to infer transmission chains, identify the source of outbreaks, and monitor the emergence and spread of new variants in near real time. These capabilities have become increasingly important for responding to emerging infectious diseases and informing public health interventions. In cancer genomics, genome assembly provides a more complete representation of tumour genomes than reference-based analyses alone, facilitating the detection of somatic mutations, large structural rearrangements, copy number alterations, and virus–host integration events that can drive tumour development [1]. This information can improve cancer diagnosis, support patient stratification, and guide the selection of targeted therapies.

Modern genome assembly algorithms typically represent reads using graph based models in which nodes correspond to reads or sequence fragments and edges represent overlaps between sequences. Two dominant paradigms are used: de Bruijn graphs and overlap graphs [34, 29]. In the de Bruijn graph approach, reads are decomposed into smaller *k*-mers (sub-sequences of length *k* base pairs) and assembly corresponds to finding an Eulerian path through the graph. Eulerian path problems can be solved in linear time, which explains the practical success of de Bruijn graph assemblers [34]. The assembler then reconstructs contiguous sequences (an uninterrupted, continuous stretch of DNA formed by overlapping smaller sequencing fragments) from the graph structure. However, the de Bruijn representation introduces other challenges such as higher sensitivity to sequencing errors and difficulty resolving long repeats in the genome sequence. The overlap graph approaches form the basis for the Overlap–Layout–Consensus (OLC) framework [29]. In this paradigm, overlaps between reads are first identified, a graph describing read relationships is constructed, and a consensus sequence is generated from a traversal of the resulting graph. In overlap graphs, reads are represented as vertices and edges correspond to suffix–prefix overlaps. Recovering the genome requires identifying a graph-consistent ordering of reads or sequence fragments. In simplified optimisation formulations, the layout stage can be cast as a Travelling Salesman Problem (TSP) [2] on the overlap graph [46]. Real sequencing data introduces additional challenges, including sequencing errors and repeated genomic regions, which can generate ambiguous or spurious connections in the assembly graph and make it difficult to identify the path corresponding to the true genome sequence [46]. These structural challenges motivate the development of advanced optimisation techniques for navigating OLC assembly graphs. In generalised formulations, key optimisation subproblems arising in genome assembly are NP-hard, and no polynomial-time algorithm is known for solving them exactly in the general case. As a result, practical assemblers rely heavily on heuristics, graph simplification, and consensus procedures. The combinatorial nature of genome assembly, therefore, motivates the exploration of alternative optimisation frameworks for navigating large solution spaces. Quantum computing has emerged as one possible approach for addressing certain classes of such optimisation problems [2, 37, 5, 44].

Despite advances in bioinformatics processing, long-read assemblers such as Canu [25] and Unicycler [53] can still exhibit important failure modes, including premature circularisation, haplotype mis-resolution, spurious reconstructed segments, and systematic read clipping that may indicate divergence between the assembly and the source reads [49]. Additionally, many long-read assemblers based on Overlap–Layout–Consensus (OLC) principles remain computationally demanding, particularly for large genomes and high repeats datasets. Classical assemblers using greedy heuristics, traversal ordering, and graph simplification may not recover globally optimal solutions [49], forcing trade-offs between computational tractability and assembly accuracy. These limitations motivate investigation of alternative optimisation strategies for de novo assembly, especially at higher scales. Quantum optimisation algorithms such as Quantum Approximate Optimisation Algorithm (QAOA) [9] and Quantum Annealing [19] aim to use superposition and tunnelling to explore complex solution spaces more efficiently than classical heuristics, potentially reducing time-to-solution for hard combinatorial subproblems in assembly in the future [37], while also offering a distribution of nearly-optimal solutions useful in identifying variations and allows room for interpretation to domain experts. In simplified formulations, parts of the assembly layout problem can be expressed as constrained path optimisation problems on a graph, making them conceptually related to the TSP. Quadratic Unconstrained Binary Optimisation (QUBO) is a mathematical framework used to represent complex combinatorial optimisation problems in a standardised form amenable to quantum/hybrid solvers [30]. Its flexibility has enabled a wide range of optimisation problems to be mapped into a common formalism. Both the TSP and genome assembly have previously been encoded in QUBO [44, 2]. While the QUBO approach for quantum genome assembly is promising, it also has limits, which causes specific challenges for practical research on real sequencing data:

- Current Noisy Intermediate-Scale Quantum (NISQ) devices have a limited number of qubits and cannot (yet) be used for practical and scalable applications [12]. NISQ devices are characterised by qubits prone to errors that lose their quantum state (decoherence) due to environmental interference. For genome assembly problems, which often require large graphs containing hundreds or thousands of vertices and edges, the limited qubit count restricts the size of assembly instances that can be encoded directly on current hardware.
- A quantum circuit is a graphical model representing a sequence of quantum gates and measurements applied to qubits to perform computations. Operating on the principles of quantum mechanics, they use gates (e.g., Hadamard, CNOT) to manipulate superpositions and entanglement. Gate fidelities of current devices for single-qubit operations are around 99 *−* 99.5% and for two-qubit gates around 95–99%, which introduce significant errors in circuits that compounds as number of operations increase [13, 40]. Genome assembly QUBOs often require deep circuits with many two-qubit interactions to represent graph connectivity and optimisation constraints. The accumulation of gate errors can therefore reduce solution quality and increase the probability of obtaining invalid or suboptimal assemblies.
- QUBO encoding can dramatically increase the number of binary variables required to represent a combinatorial optimisation problem, impacting scalability even further. This affects large scale experiments to benchmark quantum algorithms on real devices as well as ideal simulations on classical devices [47]. Overlap graphs may contain hundreds or thousands of nodes, causing the corresponding QUBO formulations to grow rapidly in size. The resulting increase in variables and interactions can exceed available hardware resources and significantly increase classical simulation times.
- QUBO is a generalized approach which works well for certain problems like Max-Cut. For problems like the TSP, there are strict constraints (e.g. each city must be visited exactly once, valid tour order, etc). Since QUBO is unconstrained, these constraints must be added as penalty terms in the objective function. These penalties only encourage the constraints rather than enforce them perfectly, so the model may produce approximate or invalid solutions if penalties are not tuned properly [47]. Similarly, genome assembly contains strict structural constraints, such as maintaining valid paths through the assembly graph and avoiding conflicting connections. Poorly selected penalty weights may lead to infeasible assemblies, disconnected paths, or biologically implausible genome reconstructions.
- The TSP formulation provides an elegant theoretical abstraction of genome assembly, but assumes that every graph node should be visited exactly once. In practical sequencing datasets, however, the assembly graph can contain redundant reads that need not all be incorporated into the final assembly, meaning that traversal of every graph node is not necessarily required for reconstruction of the genome. Graph sparsification is routinely used to remove redundant connections and reads [17]. Moreover, standard TSP-QUBO formulations largely encode graph topology while neglecting read sequences and overlap-specific information that determine biologically plausible assemblies [2, 44]. Understanding this disconnect is therefore essential for developing richer quantum compatible formulations that better capture the biological structure of genome assembly.
- Quantum and quantum-inspired algorithms such as Simulated Annealing (SA), Quantum Annealing (QA), and QAOA operate probabilistically. They are not exact solvers that guarantee the most optimal solution [9, 23, 41]. While the solvers optimize the landscape sufficiently to include valid genome reconstructions within their solution samples, identifying the correct assembly sequence from the sampled solutions remains an open challenge.

In this work, we present a quantum-assisted genome assembly pipeline for a clinically originating *Pseudomonas aeruginosa* genome using real Oxford Nanopore Technology (ONT) long reads from Ref [32]. The data set comprises real long-read sequencing data generated on an Oxford Nanopore platform. We replace the graph layout optimisation component within a Unicycler long-read assembly workflow [53], with a quantum optimisation approach (Figure 1): the assembly string graph [35] produced in Unicycler is framed as a variant of the TSP, and converted into a QUBO formulation similar to Ref [2]. To practically implement the assembly on a real device, given qubit constraints and noise of the device, we use the Hamiltonian Auto Decomposition Optimisation Framework (HADOF) [43]. HADOF is a generalised framework that can be used to solve QUBO problems on current quantum hardware at scales larger than the size of the device, by automatically federating the optimisation process into small subproblems and iteratively refining the final aggregated solution. HADOF may also improve robustness on noisy hardware by favouring smaller circuit executions, which can generally be implemented with higher fidelity on NISQ devices. We evaluate the feasibility of this approach and discuss its implications for future quantum-assisted assembly methods. The main contributions of this work are as follows.

- Integration of HADOF with QUBO formulated genome assembly to enable the solution of assembly problems that exceed the qubit and connectivity limitations of current NISQ hardware.
- Demonstration of large scale^1^ QUBO-based genome assembly on real quantum hardware enabled by HADOF, and compare its performance against a classical state of the art assembler Unicycler, as well as classical simulated annealing (SA) and ideal, noise-free quantum simulations.
- Investigation of the relationship between optimisation metrics and biological assembly quality of the sampled solutions from the probabilistic QUBO solvers using real *Pseudomonas aeruginosa* sequencing data to contrast the Unicycler and QUBO formulations, and identify future directions for practical quantum advantage in genome assembly. This analysis addresses two critical challenges: it provides empirical insights to guide the development of biologically expressive, assembly-specific QUBO models, and it establishes a data-driven selection framework using the metrics to identify correct genome reconstructions from sampled distributions without relying on a reference genome.
- An open-source implementation of HADOF and the complete quantum-assisted genome assembly benchmarking pipeline as a tool for future experiments and possibly practical usage. The data collected from previous runs, including experiments on a real quantum device, is made available on a supplementary GitHub^2^.

**Figure 1:**
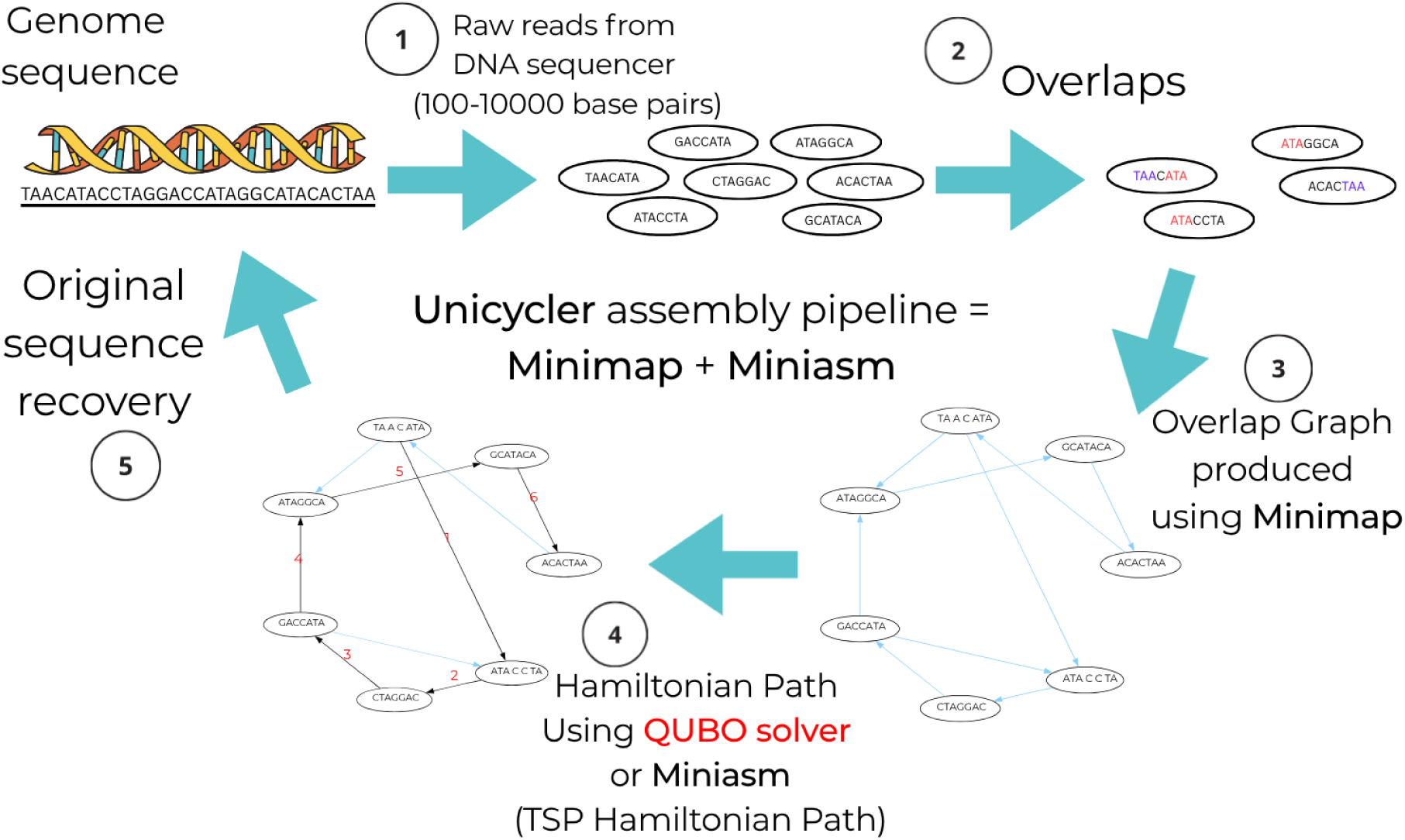
Overview of the genome assembly pipeline and the integration of the proposed QUBO approach within the Unicycler framework. (1) A DNA sequencer generates a collection of raw sequencing reads, sampled from the original genome sequence. (2) Pairwise overlaps between reads are identified using Minimap (within Unicycler), with overlapping regions highlighted to indicate shared sequence content between reads. (3) The detected overlaps are represented as a directed overlap graph, where each node corresponds to a read and an edge from read (i) to read (j) indicates that the suffix of read (i) overlaps the prefix of read (j). (4) The assembly problem is formulated as a Hamiltonian Path problem, equivalent to a TSP. Classically, it is performed using Miniasm (within Unicycler). The numbered red edges illustrate one valid traversal order through the graph. This is the NP hard optimisation step that this research attempts to replace with quantum QUBO optimisation. (5) The ordered reads are merged according to their overlaps to reconstruct the original genome sequence.

The remainder of this paper is organised as follows. Section 2 reviews existing approaches to quantum optimisation and genome assembly, highlighting the limitations that motivate the proposed framework. Section 3 describes the proposed QUBO formulation, the HADOF decomposition framework, and the assembly pipeline. Section 4 presents experimental results from *Pseudomonas aeruginosa* sequencing data and analyses the relationship between optimisation performance and assembly quality, discussing the implications for future quantum genome assembly. Finally, Section 5 concludes the paper and outlines directions for future work.

## 2 Related Work

In this section, we review the key literature in the areas relevant to our approach: classical long-read assembly methods and their limitations, QUBO and quantum-based assembly formulations, and federated/hybrid optimization frameworks. We identify critical gaps that motivate the contributions of this work and situate this work in the wider context of current quantum optimisation for genomics literature.

### 2.1 QUBO and Quantum Inspired Assembly Approaches

Classical genome assembly algorithms commonly employ graph based strategies such as the Overlap Layout Consensus (OLC) and De Bruijn Graph (DBG) frameworks. For long, noisy reads, OLC strategies build an overlap graph of reads and seek consistent traversal paths, an NP-hard optimization in general. Popular long read assemblers including Canu [25], Flye [24], Miniasm [27], and others have been benchmarked extensively on real and simulated ONT data, with performance varying significantly depending on read quality, coverage, and genome complexity [52, 26]. These benchmarks consistently demonstrate that no single assembler universally outperforms others across all metrics such as contiguity, sequence identity, and completeness, and that assembly quality degrades with high read error rates and complex repeat structure [52, 26]. Hybrid assemblers such as Unicycler, which integrate long and short reads, often recover challenging genomic elements such as plasmids more reliably than using only long reads [18]. Despite engineering progress, the underlying optimisation remains heuristic, with no guarantees of global optimality, and scaling to large, noisy datasets continues to be a major challenge.

Formulating assembly as a combinatorial optimization problem has motivated the use of Quadratic Unconstrained Binary Optimization (QUBO) and quantum computing techniques. The QUBO representation enables encoding of discrete assembly objectives, such as finding a path through an overlap graph into a binary quadratic cost function that can be solved on quantum annealers or quantum inspired algorithms. Boev et al. [2] present one of the first experimental demonstrations of QUBO-based assembly using quantum annealing on a D-Wave machine [33] on small synthetic and bacteriophage genomes. Their work embeds an OLC graph into a QUBO formulation and solves it with current quantum annealers, illustrating current hardware limitations for realistically sized problems and providing baseline benchmarks for future comparisons. Similarly, Sarkar et al. [44] implement a QUBO based assembly pipeline highlighting the algorithmic steps required to translate biological assembly tasks into quantum optimization problems. More recent work has extended the application of quantum computing in genomics beyond proof-of-concept overlap graph assembly [5, 8]. Chen et al. [5] demonstrate haplotype-resolved assembly of diploid and polyploid genomes using quantum computing techniques, highlighting the potential of quantum optimization for resolving complex genomic variation and phasing problems. In parallel, the divide-and-conquer hybrid assembly framework proposed in [8] introduces a scalable quantum algorithm for integrating short and long reads, aiming to reduce computational complexity by decomposing the assembly problem into smaller optimization subproblems. While these efforts demonstrate the feasibility and growing sophistication of QUBO and quantum-based formulations, they remain limited to relatively small graph instances or hybrid decomposition strategies that may not address the full complexity of large real-world ONT datasets at practical scales and may be harder to generalise. Existing quantum bioinformatics surveys [20] further emphasize that most quantum techniques for assembly are still at a proof-of-concept stage and that substantial methodological and hardware progress is required before clear advantages over state-of-the-art classical assemblers can be demonstrated.

### 2.2 Quantum Algorithms and Federated Optimization Frameworks

Recent research has explored improved constraint handling techniques in quantum optimisation algorithms. For example, constrained QAOA variants introduce specialised mixer gates that restrict the search space to feasible solutions [10]. Alternative approaches modify problem formulation for quantum annealing to enforce constraints directly without penalty terms [15]. Grover Adaptive Search can also incorporate constraints by flagging feasible states using additional ancilla qubits [12]. Barren plateaus in the solution space, where gradients vanish exponentially and prevent efficient optimisation of variational quantum circuits (VQCs), are a major challenge in QAOA [6]. FALQON can be used to initialise QAOA circuits and mitigate these optimisation difficulties [31]. Additional variants such as GM-QAOA and fixed-point Grover search improve convergence properties in constrained optimisation problems [36]. More recently, quasi-binary encoding schemes and hard constraint-preserving mixers have been proposed as alternatives to conventional soft-penalty formulations. Chen et al. [4] introduce a quasi-binary encoding based Quantum Alternating Operator Ansatz that preserves sum constraints directly during optimisation, avoiding the need for large penalty coefficients in constrained QUBO problems. Such approaches are particularly relevant for graph-based genome assembly formulations where incoming and outgoing edge constraints are traditionally enforced through quadratic penalties. Related studies further quantify the performance and applicability of quantum approximate optimisation algorithms in constrained combinatorial optimisation settings, including portfolio optimisation and structured QUBO problems [55].

A major challenge in applying quantum optimisation algorithms is scalability. QAOA [9] and quantum annealing [19] are typically limited by the number of qubits and connectivity available on current hardware. To address this limitation, several decomposition strategies have been proposed. Recursive QAOA (RQAOA) [3] iteratively fixes subsets of variables using QAOA solutions to reduce problem size. Other approaches decompose large graphs into smaller subgraphs that can be solved independently and then recombined [56]. Distributed optimisation frameworks extend this idea further by solving subproblems in parallel and aggregating solutions. For example, Distributed QAOA (DQAOA) [21] decomposes large QUBO instances into sub-QUBOs that can be solved across distributed quantum or classical resources. Complementary scalability strategies have also been explored through semidefinite programming (SDP) relaxations of combinatorial optimisation problems. Yuan et al. [54] demonstrate that structured QUBO and MaxCut relaxations can be solved using Pauli-sparse quantum Gibbs states and corresponding quantum-inspired classical surrogates, achieving exponential speed-ups for highly structured instances. Their work reports solving a 2^50^ variable SDP relaxation on classical hardware within 0.15% of the optimal solution, illustrating how relaxation-based methods may provide an alternative route for scaling beyond current quantum hardware limitations. These approaches demonstrate that decomposition, distributed optimisation, and relaxation-based frameworks may enable quantum algorithms to address substantially larger optimisation problems than current hardware would otherwise permit.

### 2.3 Literature Gaps and Contributions

The literature reviewed above demonstrates promising proof-of-concept studies but also reveals important gaps. Existing quantum genome assembly studies have primarily demonstrated feasibility on small synthetic datasets or very small viral genomes. Most studies rely on simulated reads, while evaluations using real sequencing data remain uncommon. We are not aware of any studies that have attempted long read genome assembly using real data on real quantum hardware. Comparative benchmarking against modern classical assembly pipelines remains underexplored, making it unclear whether quantum optimisation methods can produce biologically meaningful assemblies relative to established assemblers. Furthermore, the rapid growth of QUBO formulations with increasing graph size directly reflects the scalability challenges discussed in the introduction. Realistic assembly graphs contain hundreds or thousands of reads and overlaps, exceeding the qubit counts and connectivity available on current NISQ hardware. Existing studies therefore lack scalable decomposition strategies capable of solving large assembly graphs while preserving meaningful optimisation structure. Similarly, the approximate nature of QUBO constraint penalties raises questions regarding the relationship between optimisation quality and downstream biological assembly quality, which has received limited investigation.

The challenges discussed in Section 1 the limited qubit counts and noise of NISQ devices, the scalability limitations of QUBO formulations, identifying the most biologically plausible solutions from the optimised sample solutions and the difficulty of enforcing assembly constraints within unconstrained optimisation models - remain largely unresolved in current quantum genome assembly research. This work addresses these gaps through the contributions outlined in Section 1. We evaluate the approach on a real ONT *Pseudomonas aeruginosa* dataset, representing a genome approximately three orders of magnitude larger than previous quantum genome assembly demonstrations. Then, we employ HADOF [43, 42] to overcome current hardware limitations through decomposition, enabling execution on contemporary quantum devices. By integrating the quantum optimisation stage within a conventional overlap-layout-consensus (OLC) [29] assembly workflow similar to Unicycler, this work demonstrates compatibility with existing long-read assembly pipelines and establishes a pathway toward the practical application of quantum optimisation in genome assembly. Finally, we analyse the sampled results from the QUBO based assembly, by benchmarking against a current state-of-the-art classical assembler - Unicycler. We define metrics to evaluate the relationship between solutions arising from the abstract QUBO mathematical formulation and its corresponding genome assembly sequence to provide a principled framework for quantifying how optimisation quality translates into biological assembly quality, enabling systematic evaluation of whether improvements in the optimisation objective produce biologically meaningful outcomes. This establishes a benchmark for assessing future quantum optimisation methods beyond QUBO objective values alone.

## 3 Approach

Figure 2 represents the overall pipeline and describes the differences in the quantum and classical approaches. Raw reads data is collected from a genome sequencer and needs to be assembled to obtain the complete genome sequence of an organism. In its long-read-only workflow, Unicycler [53] invokes Minimap [28] for identfying overlaps and Miniasm [27] for assembly operations, followed by consensus-polishing and circularisation procedures. A string graph [35] representing the overlaps is created from raw sequencing reads using Minimap. This is the main input for both classical and quantum approaches. Then, Miniasm performs heuristic assembly to individually identify and resolve non-linear areas in the graph such as tips, branches, bubbles, and internal sequences. In the quantum approach, the OLC is first transformed into its QUBO TSP format. The QUBO solver then searches for a low-energy graph traversal under the encoded path constraints, rather than applying the local graph cleaning heuristics used by classical assemblers. Here, we use HADOF [43] for scalability on ibm_torino [16], a gate-based Quantum Processing Unit (QPU). Since QUBO and quantum-based optimisation are both approximate, the solutions may contain errors even after the optimisation process, but it will be significantly more organised (closer to a single linear assembly sequence) than the original string graph (the string graphs tends to have multiple incoming and outgoing edges for every node). The QUBO solvers studied in this research SA and QAOA produce distributions of approximately optimised sample solutions, instead of a single final solution like Unicycler. We post-process the results to resolve the errors that remain for all the samples. The steps after this are common to both classical and quantum pipelines. From the linear sequence of nodes (representing reads) and edges (representing overlaps) produced from the assembly step, the sequences are combined in accordance with the overlap lengths to form one contiguous sequence, called unitigs, in the unitigging step. Then, the sampled assemblies are polished for accuracy using Racon [50]. Polishing improves the accuracy of raw de novo genome assemblies by correcting errors in the unitig by aligning raw reads to the assembly, performing a partial order alignment to generate a consensus, and fixing mismatches and indels. After this step, we evaluate the final output sequences for their accuracy and compare the assemblied from the sampled QUBO solutions with the Unicycler solution using QUAST [14]. Additionally, we also perform evaluation of the raw QUBO output samples before “Cleaning Errors” or the post-processing steps. This analysis provides useful information on how to improve the QUBO formulation for genome assembly, and identifying the relationship between the sampled solutions and their corresponding assembly sequences to identify the correct solutions from the samples.

**Figure 2:**
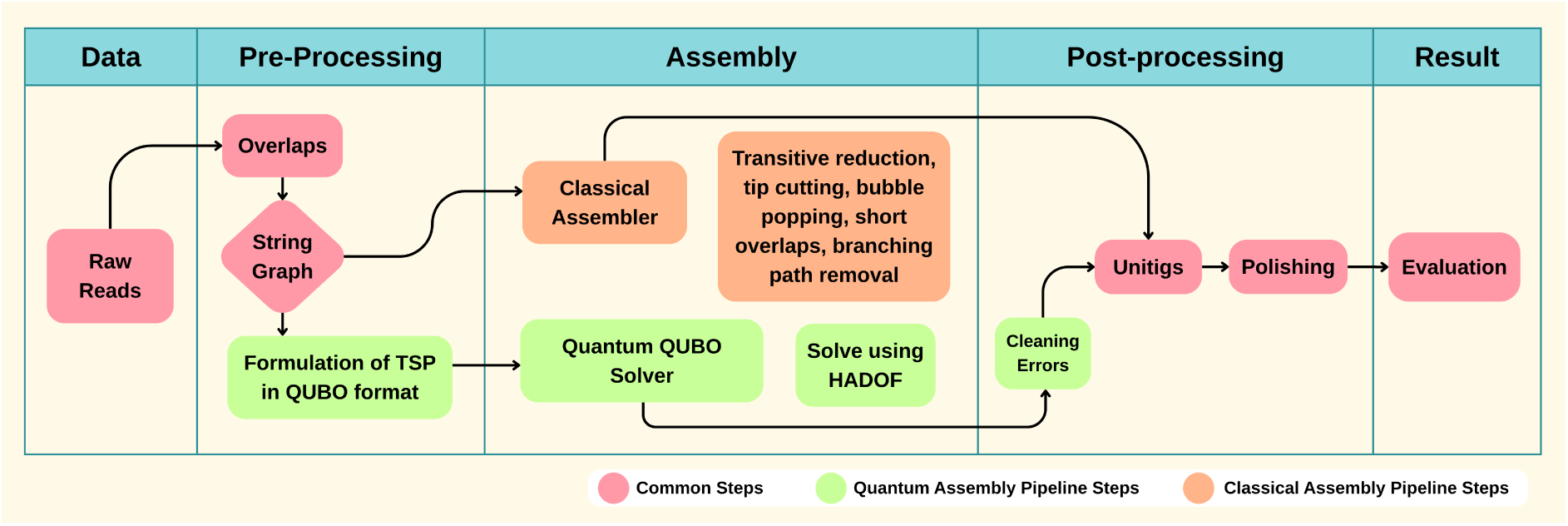
Comparison of the classical and quantum genome assembly pipelines. Overlaps are identified between raw sequencing to generate a string graph, which forms the common input to both approaches. The classical pipeline uses graph simplification heuristics, including transitive reduction, tip cutting, bubble popping, short-overlap filtering, and branching path removal to generate assembly paths. The quantum pipeline reformulates the assembly problem as a TSP in QUBO form and solves it using HADOF. The quantum approach requires a post-processing step to produce a final linear path. Both approaches subsequently perform unitig generation, polishing, and assembly evaluation. Pink blocks denote steps common to both pipelines, orange blocks indicate classical assembly operations, and green blocks represent quantum specific stages.

### 3.1 Data

ONT long-read sequencing data from *Pseudomonas aeruginosa* [32] was used to evaluate the proposed approach on a real genome assembly problem. The dataset consists of 21,969 Oxford Nanopore Technology (ONT) reads obtained directly from a DNA sequencer. The reads range from 1 bp to 477,150 bp in length, with an average read length of 10,991.7 bp. The datasets and source code is available on the companion GitHub^3^ repository. This dataset was selected because it represents a realistic long-read assembly problem containing sequencing errors, ambiguous overlaps, and graph complexities that are encountered in practical genome assembly, at a scale 1000x larger than current quantum genome assembly problems addressed in the literature (the largest genome assembled using a quantum pipeline, as far as we are aware of, is around 5 kilo bp using a simulated dataset of the *ϕ*X 174 Bacteriophage by Boev et al. [2]).

### 3.2 Pre-Processing

Within Unicycler, Minimap constructs a string graph [35] for long-reads by first computing all-versus-all overlaps between raw reads and then passing these alignments to Miniasm, a dedicated assembly tool that builds and simplifies the graph. The quantum assembly pipeline formulates a QUBO from the string graph which is optimised using a quantum algorithm.

#### 3.2.1 Classical Steps

Minimap identifies exact k-mer^4^ matches between reads and anchors them into linear chains to form stable local alignments. These chains allow Minimap to specifically isolate suffix-prefix relationships where the terminal sequence of one read aligns with the start of another. Finally, these detected overlaps are exported in Pairwise Alignment Format (PAF), providing a comprehensive record of read-to-read alignments and their respective overlap lengths.

The string graph method [35] is commonly used in OLC based assemblers. Redundant overlaps are removed using transitive reduction, and linear paths are collapsed, producing a compact graph that preserves true read relationships. Miniasm takes the following steps to complete the assembly: It uses the ONT reads sequences in the FAST-All (FASTA) format and the overlaps from Minimap in the Pairwise mApping Format (PAF) to produce a raw string graph in the Graphical Fragment Assembly (GFA) format [35]. The GFA is a graphical format with edges connecting overlapping nodes representing the reads which are potentially continuous sequences. This graph is used to create the QUBO formulation of the assembly problem as shown in Figure 2. The resulting graph from *Pseudomonas aeruginosa* contained 524 nodes and 2,313 directed overlap edges. The edge-variable QUBO formulation therefore contained 2,313 binary variables. A direct one variable per logical qubit implementation would require 17.4 times the 133 physical qubits available on ibm_torino; HADOF instead executes this through five-variable decompositions. It uses 462 5-qubit circuits and one 3-qubit circuit to accommodate for a total of 2,313 variables since it is not divisible by 5.

#### 3.2.2 QUBO Formulation

Along the lines of Ref. [2], we formulate the problem of finding a Hamiltonian path in the string graph as a QUBO problem. Let a directed OLC graph be given by *G* = (*V, E*), where *V* = *{*1, 2*, …, N}* is the set of vertices and *E* is the set of directed edges (*u, v*), with *u, v ∈ V*. In the standard permutation-matrix formulation, the Hamiltonian path is represented by an *N × N* binary matrix *X* = (*x_v,i_*), where *x_v,i_* = 1 indicates that vertex *v* is visited at position *i* in the path. This representation requires *N*^2^ logical binary variables for an *N* -vertex graph. The corresponding QUBO is

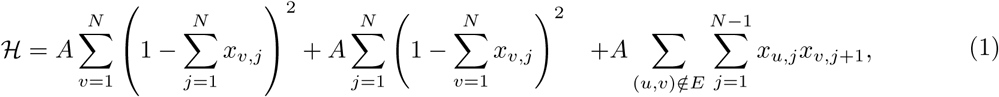

where *A >* 0 is a penalty coefficient. The first two terms enforce that each vertex is visited exactly once and that each position in the path contains exactly one vertex, respectively, while the third term penalises transitions between vertices that are not connected by a directed edge in the string graph [2].

For sparse, predominantly acyclic string graphs, the number of binary variables can be reduced by associating a binary variable directly with each graph edge. Let

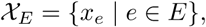

where *x_e_* = 1 indicates that edge *e* is selected as part of the assembly. The number of logical variables is then

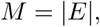

which can be substantially smaller than the *N*^2^ variables required by the permutation-matrix representation.

In our implementation, the edge variables are used to construct a QUBO directly from the local connectivity of the directed string graph. For each vertex *u ∈ V*, the incident edges are separated according to their direction relative to the vertex:

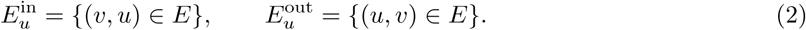

A negative diagonal coefficient is assigned to each edge at each of its endpoints, providing a negative linear contribution that favours the selection of edges. In addition, pairs of edges that are both incoming to the same vertex or both outgoing from the same vertex receive a positive quadratic interaction. The resulting QUBO objective is

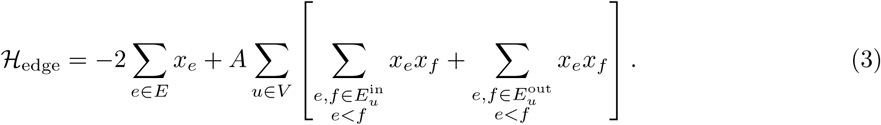

In this formulation *x_e_* = 1 represents the inclusion of an edge and *x_e_* = 0 represents its exclusion. The negative linear term rewards the inclusion of graph edges, whereas the quadratic terms penalise branching by increasing the objective when multiple selected edges enter or leave the same vertex. An incoming and an outgoing edge incident on the same vertex do not generate a quadratic penalty. Consequently, the formulation favours configurations in which selected vertices are connected by a single incoming and outgoing edge, while discouraging configurations containing multiple incoming or outgoing edges at the same vertex. The resulting solutions therefore tend toward non-branching path- or cycle-like structures. Unlike the permutation-matrix formulation, however, the edge-based QUBO does not explicitly enforce that every vertex is visited exactly once and should therefore be regarded as a relaxed connectivity formulation rather than a strict Hamiltonian-path encoding. In the experiments, the interaction coefficient was set to *A* = 2, providing a quadratic penalty for competing incoming or outgoing edges while retaining a negative linear reward for edge selection. This choice discourages branching without imposing a hard degree constraint, allowing the optimisation to balance edge inclusion against local connectivity conflicts. The edge-based formulation also does not explicitly encode the position of a vertex in the traversal.

Instead, vertex ordering is represented implicitly through the directed edges selected by the optimisation. This reduces the number of logical variables from *N*^2^ to *M* = *|E|*, providing a substantial reduction for sparse string graphs. Such a reduction is particularly advantageous for near-term quantum devices, where limited qubit counts, restricted qubit connectivity, and accumulated gate errors constrain the size of QUBO problems that can be embedded and solved. The QUBO solvers in this research produce a distribution of sample solutions

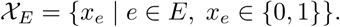

### 3.3 Assembly

In the classical Unicycler assembly method, Miniasm performs the assembly of the string graph into a contiguous sequence. The assembly is performed using heuristic steps of transitive reduction, tip cutting, bubble popping, short overlap cutting, internal sequence removal, one more round of short overlap cutting, and a final step of branching path removal to produce the final string graph [27, 53].

#### 3.3.1 Classical Assembly Steps

##### Transitive Reduction

If read A overlaps B, B overlaps C, and A also overlaps C, the edge from A to C is transitive (redundant) because the path through B already provides that information. Miniasm removes these transitive edges. This drastically reduces the number of edges without losing any connectivity information, turning the “hairball” graph into a more linear structure.

##### Tip Trimming

Tips are dead-end paths in the graph caused by reads with sequencing errors at their ends or missing overlaps. Miniasm identifies nodes with an out-degree of 0 (no outgoing connections) and trims them if they are composed of only a few reads (the default is 4 or fewer).

##### Bubble Popping

Bubbles occur when there are two or more different paths between two nodes. In long read data, these are usually caused by small errors or variations between different copies of a chromosome. Bubbles are popped by picking one optimal path and removing the others, which collapses the bubble into a single linear path.

##### Short Overlap Cutting

In some cases, reads might overlap by a very small number of bases due to random chance or because of repetitive sequences. Overlaps that fall below a specific length or quality threshold are discarded. This breaks weak, false connections that would otherwise create tangles in the graph.

##### Internal Sequence Removal

Sometimes a read is entirely contained within another longer read, or a sequence is redundant because it is already represented by a better path. Miniasm identifies and deletes nodes that do not contribute new genomic information. The graph becomes more concise, focusing only on the unique sequences.

##### Second Round of Short Overlap Cutting

After removing internal sequences and popping bubbles, new weak connections often become visible that were previously hidden by the complexity of the graph. A second pass of the overlap filter is performed. This ensures that the remaining structure is supported only by the most robust evidence.

##### Branching Path Removal

At this final stage, if the graph still has ambiguous forks in the path that couldn’t be resolved by the previous steps (often due to unresolved repeats), the assembler uses specific logic to trim them. It removes edges that lead to ambiguous branching if they are significantly lower in coverage^5^ than the main path. This produces the final, cleanest version of the string graph, ideally resulting in circularized chromosomes or large, linear contigs.

#### 3.3.2 Quantum Optimisation for Assembly using HADOF

Section 3.2.2, explains the QUBO formulation from the string graph. This section explores the solution of the QUBO using quantum algorithms. Due to qubit and connectivity limitations of current quantum devices, the full QUBO as formulated above cannot be solved directly on existing devices [2]. We employ HADOF [43], which decomposes the global QUBO into subproblems solved iteratively on quantum hardware or simulators, and then aggregated into a global solution distribution, as shown in Figure 3. This enables scalable quantum optimization beyond native device limits. HADOF can be used flexibly with the different quantum optimisation methods, and here we use trotterized quantum annealing (QAOA-t) [41], a variation of QAOA, which is compatible with gate based devices and benchmark it against classical SA. SA was selected as the primary QUBO baseline because it optimises the same QUBO formulation without decomposition and is as a strong classical heuristic for combinatorial optimisation problems [22]. Comparing HADOF against SA therefore isolates the effects of decomposition and quantum execution while avoiding confounding factors introduced by differences in assembly algorithms or heuristics.

**Figure 3:**
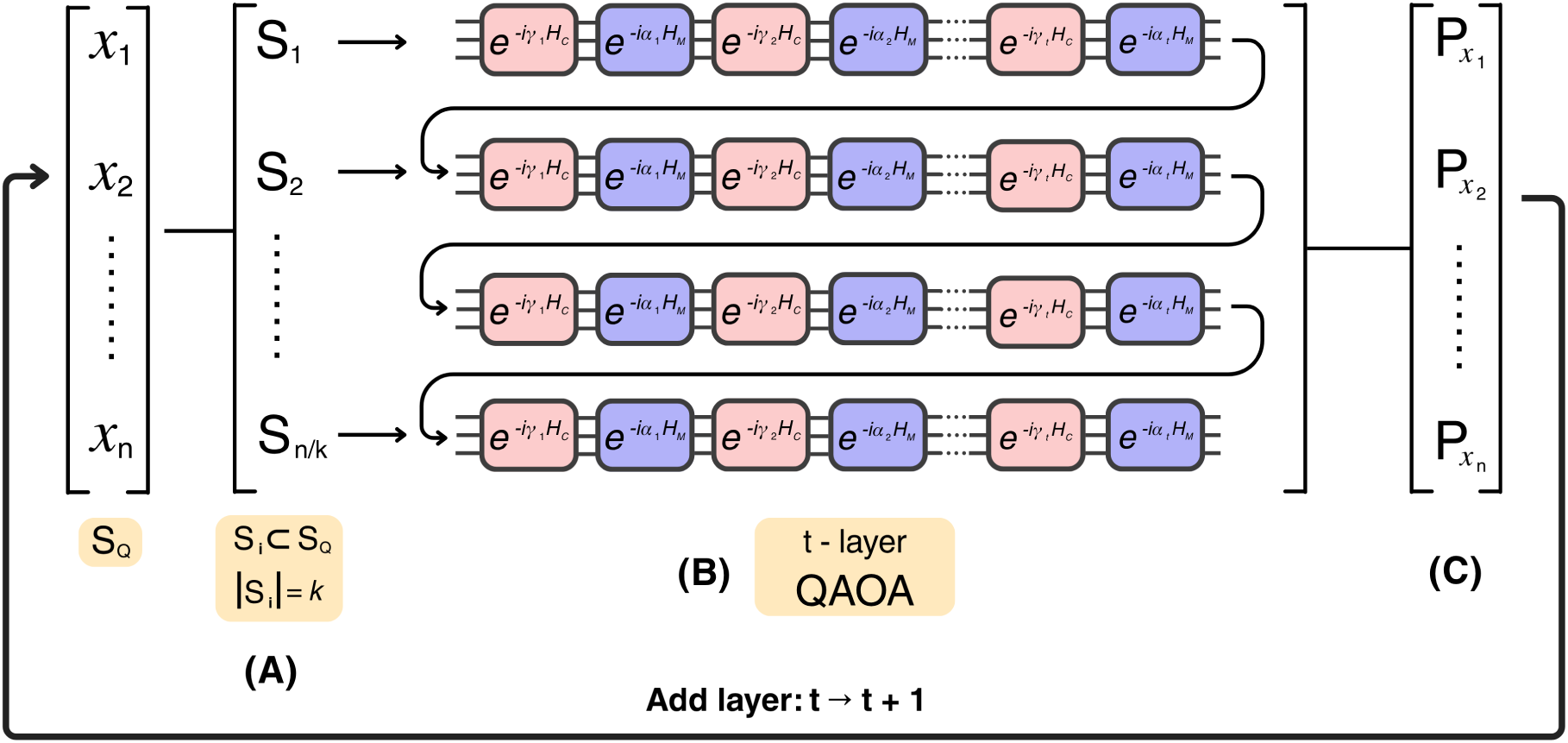
General overview of HADOF. Here, we use QAOA as the optimiser, which is called iteratively. **(A)** We choose subsets of sizes 5 from the binary variables of the global problem. These are used to form the sub-Hamiltonians *S_i_* using the expected value of each qubit from the previous iteration *P* (*x_i_*). **(B)** The QAOA circuit set up with *t* = 1 layers and in every iteration we add a layer. In this study, we use up to 5 layers, implying 5 iterations for each sub-Hamiltonian. The QAOA optimises the 5 qubit sub-Hamiltonian problem. **(C)** Once the *n/k* sub-Hamiltonians are optimised, we sample and aggregate them to form the global solution probability distribution. The single qubit expectation values *P* (*x_i_*) are used in the next iteration, while the samples aggregated form individual solutions for the QUBO problem.

The Hamiltonian Auto Decomposition Optimisation Framework (HADOF) is a procedure for approximately solving large QUBO instances that cannot be embedded directly onto current quantum hardware. We assume a global QUBO objective

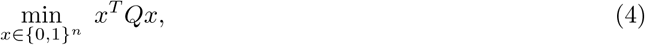

with binary decision variables *x* = (*x*_1_*, …, x_n_*), and we maintain a global “belief” vector *P ∈* [0, 1]*^n^* where *P* (*x_j_*) = E[*x_j_*] is the current estimate of the marginal probability that variable *x_j_* is 1.

##### Subproblem construction

Figure 3 (A) shows at each global iteration we select a collection of subsets *S_i_ ⊂ {*1*, …, n}* with fixed size *k ≪ n*, e.g., random or graph-informed blocks. Larger subcircuits require less QPU usage time, as the decomposed subproblems will be larger, resulting in fewer HADOF iterations [42, 43]. However, in these experiments 5 qubit circuits are used as they showed better accuracy using HADOF on NISQ devices. This is because smaller circuits are easier to embed and experience less impact from noise on NISQ devices [13].

For a given subset *S_i_*, the remaining variables are *S̅_i_* = *{*1*, …, n}* \ *S_i_*. We construct a sub-QUBO from the original obective, 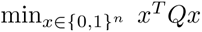, using only *x_S_* by replacing each inactive variable *x_j_* (for *j ∈ S̅_i_*) by its expected value *P* (*x_j_*), i.e,

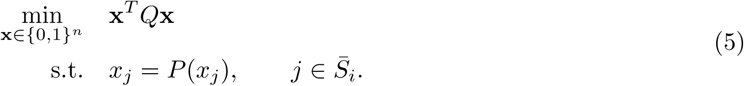

which yields a *k*-variable QUBO (and therefore a *k*-qubit Ising Hamiltonian after the standard *{*0, 1*} ↔ {±*1*}* mapping [41]). This step injects “global context” into each subproblem while keeping the quantum optimisation dimension fixed. For the first iteration of HADOF, we start with an unbiased expected value for all inactive variables *P* (*x_j_*) = 0.5.

##### Quantum optimisation and marginal update

Figure 3 (B) shows QAOA optimisation. Each sub-QUBO *H_i_* is optimised using a sampler that returns a distribution over bitstrings on the *k* active variables. We use QAOA implemented with an annealing-style parametrisation, so that at depth *l* the parameters 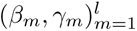 follow a fixed schedule [41] (rather than being classically trained). For each subproblem we execute the depth-*l* circuit, collect measurement samples of each qubit, and estimate per-variable expectations on the active set,

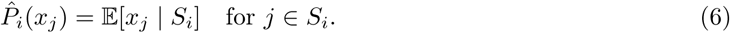

We then merge these local estimates into the global probability vector *P*.

##### Iterative refinement

Figure 3 (C) shows the algorithm proceeding for *p* global iterations. We start with an uninformative prior *P* ^(0)^(*x_j_*) = 0.5 (but this could also be a problem-informed initialisation) and gradually increase the QAOA depth from *l* = 1 to *l* = *p*, which mirrors a coarse-to-fine optimisation strategy: early iterations provide inexpensive, noisy marginals, while later iterations concentrate probability mass around higher-quality assignments. In the final iteration, we sample all the solutions from each sub-QUBO circuit and aggregate them to form solutions.

After each sweep of the *n/k* circuits, we add one layer. To measure the individual qubits to update *P* (*x_i_*) between iterations, we use 500 shots per qubit measurement and then use the average as the updated *P* (*x_i_*). In the final iteration, each circuit is sampled over all *k* qubits (instead of sampling individual qubits) using 5000 shots per circuit, to produce a distribution over each sub-solution (these are not individual qubit samples of 1 and 0, but 5-string samples, e.g, 01011 for 5 qubit circuits). 5,000 global solutions are formed by concatenating sampled sub-solutions in sampling order, to produce global solution strings of length equivalent to the number of binary variables, which can be evaluated for their objective values. For the experiments in this research, all the sub-QUBOs were of size 5 embedded on 5 qubit circuits. Previous experiments [42] show stable accuracy while scaling up problem size using 5 qubit HADOF decomposition.

HADOF converts a single large QUBO into a sequence of small, hardware-feasible sub-QUBOs, and uses the marginals extracted from quantum samples to propagate information between subproblems. The output of HADOF is a set of sampled global candidate solutions. *P* (*x_i_*) = 0.5 is used as initialisation for all *i*. Circuits use 5 layers with *β_m_* = 1 *−* (*m/*5) and *γ_m_* = *m/*5. Increasing the number of layers in the circuit improves the theoretical accuracy of the optimisation procedure as the circuit approaches the continuous adiabatic limit, in ideal noise-free settings. However, in practical physical implementations, accuracy eventually declines due to hardware constraints and noise [39].

### 3.4 Post-processing and Assembly Refinement

#### 3.4.1 Classical Post-Processing Steps

In the classical Unicycler pipeline, after the string graph has been simplified using the heuristics (like bubble popping and tip cutting as noted in Section 3.3), the process moves from a graph of connections to a linear sequence of DNA. To form a unitig (contiguous sequences from the assembly path) from a linear graph, the assembler takes the sequences of every pair of overlapping reads from the final assembled graph. Since we know the exact overlap length between the reads from the previous mapping steps, the assembler stitches them together, discarding the redundant overlapping portion of the second read. Because Miniasm uses the raw sequences of the reads, the resulting unitig inherits the high error rate of the original ONT reads. This sequence is technically contiguous but contains many noisy bases, insertions, and deletions, which are resolved in a polishing step by using Rapid Consensus (Racon) [50]. Polishing is the process of transforming a draft assembly into a high accuracy consensus sequence. The raw sequencing reads data used to build the assembly are mapped back onto the draft unitigs. Even if the unitig has an error at a specific position, the majority of the raw reads might show the correct base. Instead of just looking at one read at a time, the polisher uses Partial Order Alignment. It creates a mini-graph of all the reads covering a specific segment of the unitig. By looking at the pileup of reads, the algorithm determines the most likely base at every position. The most common error in ONT data are indels (Insertions/Deletions, e.g., a string of five As being read as four). Polishing adds or removes these bases to match the statistical consensus of the raw reads.

#### 3.4.2 QUBO Pipeline Post-Processing Steps

An additional post-processing step is required for the quantum assembly before the unitigging step. The output from the QUBO solver gives us a graph which is a subset of the string graph. The penalties are designed such that the optimisation attempts to maximize the number of reads (nodes) which have exactly one node before it (input edge) and one node after it (output edge). However, the QUBO solvers are approximate [13, 40] and significant errors arise from HADOF during decomposition and real device noise. Our evaluation first attempts to analyse the errors arising from different QUBO solvers in the graphical stage. Then, we use classical postprocessing to obtain the longest linear chain in the graphical solution to produce a final sequence, which can be evaluated using QUAST [14], a quality assessment tool for genome assemblies.

During the post-processing, the nodes violating the constraint above are identified. Initially, nodes which have an in-degree or out-degree more than 1 are resolved. A Depth First Search (DFS) [48] is used to find the in-edge and out-edge with the longest upstream and downstream paths, and remove all excessive edges. For nodes that have an in-degree or out-degree less than 1, all possible reconnecting edges are identified merged using DFS to reconnect the edge with the longest upstream or downstream path. We finally choose the longest continuous path as the final assembly and write this solution as a GFA file.

GFAtools [38] is used to produce unitigs (a singular sequence of base pairs) from the GFA. GFAtools does not perform any changes to the path produced from the nodes of the string graph, since it is already linear and assembled, it only merges the strings based on the overlap of the sequences. This output is converted into the FASTA file format and then polished for 4 rounds using Racon [50] to produce the final assembly sequence, in the same way classical Unicycler does. Bacterial genomes, such as *Pseudomonas aeruginosa*, are circular in nature [32]. An additional post-processing step is required for these genomes to close linear sequences into continuous loops to match the true biological shape. Without it, assemblers mistakenly generate duplicate end sequences that break genes in half and ruin downstream genomic analyses. Unicycler solves this natively by tracking loops inside an assembly graph and using long reads to confidently resolve repetitive paths. It then seamlessly trims away the redundant overlapping ends. To remove potential terminal redundancy, in the QUBO pipeline, a reference-free terminal-overlap trimming procedure was applied. The first and last 700 kb of each assembly were extracted and aligned against one another using Minimap2 with the ‘asm5’ preset [28]. A terminal overlap was considered significant when it was at least 10 kb in length, had a minimum sequence identity of 99%, and reached within 2 kb of the corresponding sequence termini. When such an overlap was detected, the longest qualifying terminal overlap was removed from one end of the assembly to eliminate the duplicated sequence. Assemblies without a qualifying terminal overlap were retained unchanged.

### 3.5 Summary of Differences in Classical and Quantum Approaches

The fundamental distinction between the classical Unicycler approach and the proposed quantum framework lies in how they resolve the complexity of the string graph heuristic vs. holistic resolution. Classical Unicycler relies on a series of local heuristic cleaning steps (e.g., bubble popping, tip cutting) to simplify the graph incrementally until a linear path remains. In contrast, the quantum approach treats the assembly as a global optimization problem. By encoding the graph into a QUBO format, the solver attempts to find a single global minimum that represents the optimal Hamiltonian path through the entire graph simultaneously. Where the classical pipeline explicitly removes erroneous edges based on rules, the quantum solver suppresses suboptimal paths through energy penalties in the Hamiltonian. The classical pipeline is deterministic; the quantum pipeline is probabilistic and approximate, requiring a specific post-processing phase (via DFS) to fix violations of the Hamiltonian constraints caused by NISQ device noise or HADOF decomposition approximations.

Implementing genome assembly on current quantum hardware requires further navigating the limitations of the Noisy Intermediate-Scale Quantum (NISQ) devices, specifically qubit counts and connectivity. The string graph, for even small genomes, generates a QUBO with thousands of variables at least. Direct mapping to a gate-based device is often impossible due to the limited number of qubits on current devices and limited connectivity between these qubits, required by the quadratic terms in the QUBO. Each pair of variables in quadratic terms requires a connection in the device between the corresponding qubits that represent them. However, current devices rarely offer a layout where every qubit can be connected to every other variable. In these cases, transpilers [16] are used, that transform abstract, high-level quantum circuits into optimized, hardware-compatible, low-level circuits. Transpilers overcome disconnected qubits by mapping frequently interacting qubits to neighboring physical positions and inserting SWAP gates to move quantum information across the chip. This rerouting allows non-adjacent qubits to interact, though it increases the circuit’s length and susceptibility to noise.

## 4 Results and Discussion

We analyse samples of QUBO solutions for the assembly of real sequencing data from the *Pseudomonas aeruginosa* genome [32] using SA, HADOF-based QAOA simulated on a classical computer without quantum device errors (HADOFQAOA Ideal), and HADOF-based QAOA on the ibm_torino Quantum Processing Unit (QPU) [16]. A major hindrance to the practical applicability of quantum optimisation is that current quantum devices are small in qubit count and error prone [13, 40]. Many studies implementing genome assembly on real devices are limited to toy problems or manual pre-processing and quantum-classical hybrid optimisation methods [2, 44].

All the classical simulations for this research were run on Macbook Pro M3 using Qiskit [16]. Ideal quantum simulations were performed on the Aer Simulater. ibm_torino was used for real device experiments. On this Qiskit platform, the scheduler automatically transpiles the circuits. ibm_torino is a 133-qubit Heron processor featuring a hexagonal, heavy-hexagonal, or lattice-based connectivity layout as used in February, 2026 [16]. Simulated Annealing (SA) from D-Wave samplers [7] was used as a benchmark for comparison against a classical QUBO solver. SA used the sample_qubo function with anneal_time = 20. The raw sequencing data is first converted to the string graph GFA using Minimap from Unicycler. This is the first input for the QUBO assembly pipeline. The string graph consists of 524 nodes and 2,313 directed overlap edges, as in Figure 4 (A). An important feature to be noted in the string graph of this genome is that it contains multiple cyclical paths. Although the OLC based QUBO assumes the graph to be acyclic [2], the string graph from real data may contain cyclical paths. For specific genomes, such as principal bacterial chromosomes, the topology is expected to be circular [32]. The Unicycler assembly also produced a circularised chromosomal assembly. The solution from Unicycler is used as a reference for comparison.

**Figure 4:**
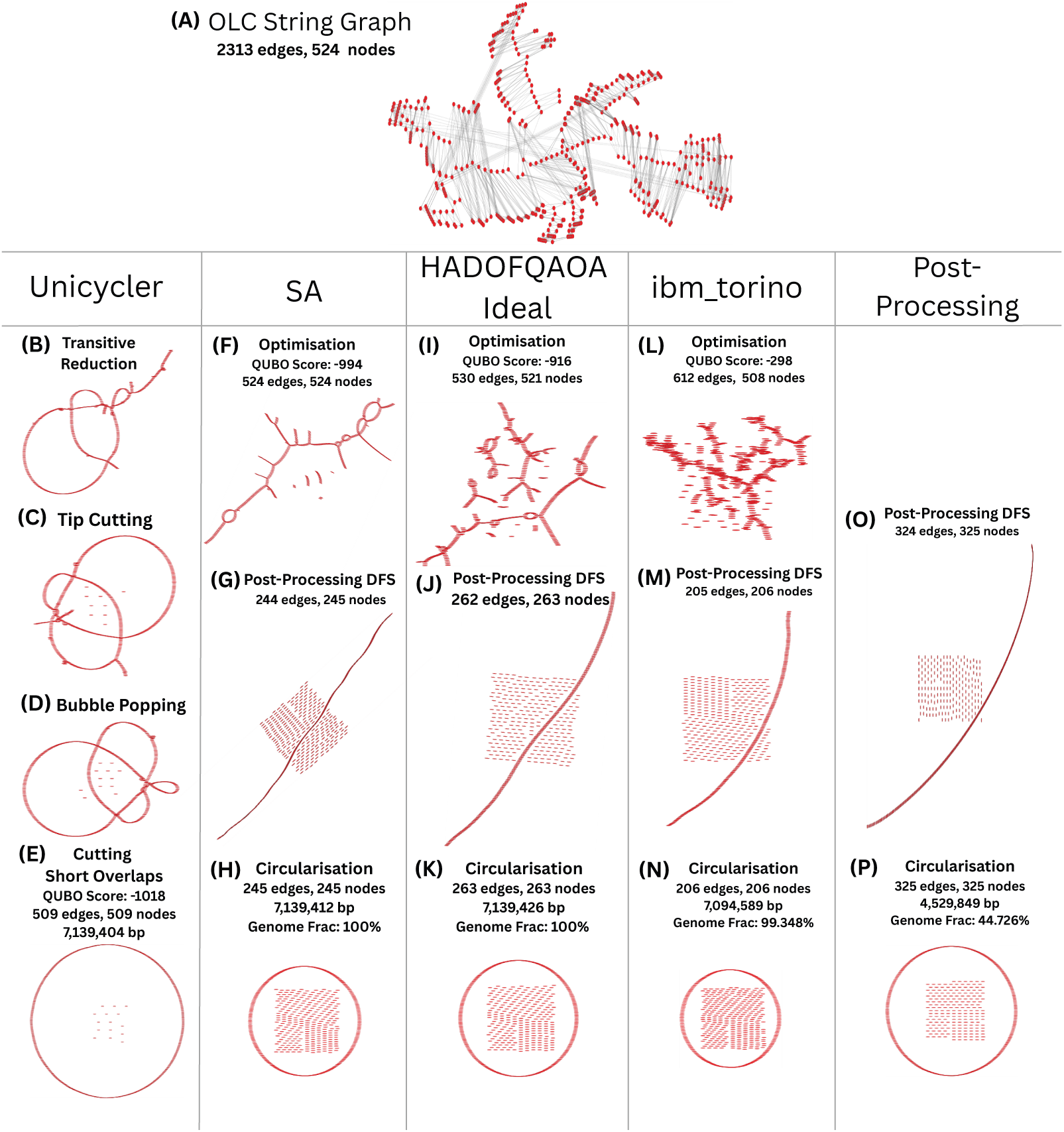
Comparing the OLC string-graph assembly using Unicycler and QUBO optimisation pipelines. Representative solutions from SA, HADOFQAOA Ideal, and ibm_torino are compared in terms of QUBO score, graph structure, assembly length, and genome fraction. Disconnected nodes in the middle of figures represent the nodes eliminated from the final assembly. The last column compares the isolated contribution of post-processing without the QUBO input (directly applying DFS on the string graph). The result implies that for this example, DFS post-processing alone produces a suboptimal assembly, and the QUBO input is essential for the generation of good assemblies.

We evaluate the QUBO-based genome assembly sample solutions at two complementary levels: the quality of the sample solutions produced by each solver and the biological quality of the genome sequences reconstructed from those solutions. To assess these two levels consistently, we first define a set of optimisation and assembly metrics in Section 4.1. The optimisation metrics here capturing the quality and structure of the QUBO solutions are the QUBO objective value, number of nodes retained in the final path and final sequence length. The assembly metrics, from QUAST [14], quantify the resulting biological reconstruction through genome fraction and duplication ratio in comparison to the Unicycler reference solution. In this evaluation we try to characterise the distribution of the sample solutions to see how well the solvers performed on the formulated QUBO, understand the relationship between the formulated QUBO and the actual objective of genome assembly, and provide insight on how to identify biologically plausible solutions from the sampled solutions. Together, these metrics provide complementary measures of whether a mathematically favourable QUBO solution also produces a biologically meaningful genome assembly. Section 4.2 presents the chosen sample solutions from Figure 4 for each solver. These solutions are chosen from the sample solutions as they exhibit biologically plausible genome reconstructions as compared to the baseline Unicycler solution.

Using the metrics defined, we first characterise the optimisation behaviour of the different solvers in Section 4.3, establishing how effectively each approach solves the underlying QUBO formulation independently of downstream sequence quality. We then examine the relationship between the optimisation and assembly metrics in Section 4.4, allowing us to determine whether improvements in the QUBO objective or graph traversal characteristics translate into improvements in genome reconstruction. Finally, Section 4.4 also provides a discussion on identifying biologically plausible assemblies from the sampled solutions. This layered evaluation is important because the QUBO objective represents an abstract optimisation criterion, whereas genome fraction and duplication ratio capture the actual biological reconstruction quality. Comparing these levels therefore reveals the performance of the current formulation for genome assembly quality, and provides insight on how to improve the QUBO formulation for assembly-specific tasks.

### 4.1 Optimisation Metrics and QUAST Metrics

To evaluate biological assembly quality, optimisation metrics of the QUBO formulation are analysed alongside the metrics from QUAST [14]. The optimisation metrics used are QUBO score, number of nodes in the final path and total assembled genome length. The metrics from QUAST are genome fraction and duplication ratio.

**QUBO score** is the objective value minimised during optimisation, representing the quality of the selected graph traversal under the QUBO formulation.

**Number of nodes in the final path** are the graph nodes retained in the reconstructed assembly path after post-processing.

The **total assembled genome length** represents the cumulative size of the final assembly in base pairs and acts as an indicator of how much genomic sequence has been reconstructed. Larger assembly lengths generally correspond to more complete traversals of the overlap graph, although excessively large assemblies may indicate repeat-induced over-assembly or incorrect graph traversal.

The optimisation metrics above are available to us without the need for a reference genome. These metrics could be helpful to identify biologically plausible assemblies from the sample solutions, if a relationship could be established between the optimisation metrics and the QUAST metrics. The QUAST metrics compare the sampled assembly solutions against the Unicycler assembly used as a reference for evaluation of accuracy.

The **genome fraction** measures the percentage of the reference genome covered by the assembled contigs. We compare how much of the assembly sequence from Unicycler was captured by the QUBO assemblies sampled. This metric provides a direct biological measure of assembly completeness, where higher values indicate that a larger proportion of the true genome has been reconstructed.

The **duplication ratio** measures the proportion of the assembled sequence that is aligned to the reference genome more than once, indicating redundant or duplicated sequence in the assembly. A higher duplication ratio generally suggests that some genomic regions have been assembled multiple times, potentially due to repetitive sequences, unresolved repeats, or assembly errors. A duplication ratio of 1.0 implies that the amount of aligned sequence is approximately equal to the reference coverage, indicating an accurate assembly.

### 4.2 Sample Assembly Results from QUBO Optimisation

Figure 4 (B)-(E) outline the steps, taken by Unicycler and present the resulting graphs after each step. The figure shows only 4 steps. The remaining steps mentioned in Section 3.3 internal sequence removal, second round of short overlap cutting and branching path removal are not shown in the figure, as Unicycler finished this assembly of the OLC within the first 4 steps. The final Unicycler assembly consists of 509 nodes as in Figure 4 (E), leaving 15 unused nodes for building the final assembly sequence.

The resulting assemblies obtained from different QUBO solving strategies are compared against the assembly generated by Unicycler, which serves as a classical baseline. Figure 4 shows sample solutions from the different solvers. The Unicycler assembly produced a circularised genome with 509 nodes, a total assembly length of 7,139,404 bp, and a QUBO objective score of *−*1018, establishing an empirical reference point for biologically valid solutions. The theoretical minimum QUBO score for the graph is *−*1048, corresponding to the maximum possible inclusion of graph edges and nodes under the optimisation formulation. However, the Unicycler result demonstrates that the biologically optimal assembly does not necessarily correspond to the absolute theoretical minimum. Instead, a subset of nodes and edges may be intentionally excluded to remove ambiguities, repeats, or low-confidence regions while still yielding a valid circular genome.

Figure 4 (F)-(N) visually presents corresponding graph transformations for a single representative QUBO solutions obtained using simulated annealing (SA), HADOF simulated on a classical machine in noise-free scenario (Ideal HADOF), and the ibm_torino quantum processor. The presented solutions were selected to have good genome fraction and similar length to the reference genome. For each solver, the graph obtained directly from the QUBO optimisation is first subjected to the DFS post-processing and circularisation before downstream polishing steps. The SA solution initially selects 524 edges and 524 nodes, with a QUBO score of *−*994. DFS substantially reduces the graph to 244 edges and 245 nodes, with a corresponding QUBO score of *−*448. Following circularisation, the resulting assembly contains 245 nodes and has a total length of 7,139,412 bp, achieving 100% genome fraction. This demonstrates that a QUBO solution with a substantially higher objective value than the theoretical minimum can nevertheless produce an assembly with comparable length and complete genome coverage.

The HADOFQAOA ideal solution follows a similar progression. The initial optimisation produces a graph containing 530 edges and 521 nodes, with a QUBO score of *−*916. Post-processing reduces this to 262 edges and 263 nodes, with a QUBO score of *−*524. After circularisation, the resulting assembly contains 263 nodes and has a total length of 7,139,426 bp, achieving a genome fraction of 100%. Although the HADOF solution contains substantially fewer nodes than the Unicycler baseline, its final assembly length remain very close to the classical reference with 100% genome fraction, demonstrating that the HADOF optimisation can identify a compact subset of the string graph that preserves the majority of the genome.

The solution obtained on the ibm_torino processor exhibits a more fragmented optimisation result. The initial QUBO optimisation selects 612 edges across 508 nodes and obtains a QUBO score of *−*298. Post-processing reduces this to 205 edges and 206 nodes, with a QUBO score of *−*410. Circularisation subsequently produces an assembly containing 206 nodes and a total length of 7,094,589 bp, corresponding to a genome fraction of 99.348%. Despite the substantially higher QUBO objective value and the smaller number of retained nodes, the resulting assembly remains close to the expected genome size and recovers more than 99% of the reference genome. These results illustrate that all three QUBO solvers managed to reconstruct biologically plausible assemblies similar in quality to Unicycler in the sampled solutions. However, the QUBO objective score is not effective to identify good quality assemblies: the structure and connectivity of the selected nodes and edges, together with the subsequent graph post-processing, play an important role in determining the quality of the final assembly.

Figure 4 (O) and (N) show the isolated effect of DFS post-processing and circularisation directly on the string graph, skipping the QUBO optimisation. Since this step is deterministic, it only produces one solution. Although the DFS manages to preserve more nodes (325 nodes), the final assembly is only 4,528,849 bp long and covers only 44.726% of the reference genome. This implies that the additional post-processing step cannot alone produce a good assembly and a good QUBO optimised input is an essential step to produce a biologically plausible assembly for this genome.

### 4.3 Performance of the QUBO Solvers

Across all experiments, SA consistently achieved the lowest QUBO objective values, frequently approaching the Unicycler reference score of *−*1018 (Figure 5 (A)). The theoretical minimum score for this assembly based on the QUBO formulation is *−*1048. The minimum QUBO score from SA is *−*1008, with an average score of *−*992. HADOFQAOA ideal simulations also produced competitive optimisation results averaging around *−*906, whereas executions on ibm_torino produced substantially higher objective values due to current hardware noise and sampling limitations, averaging around *−*370. Beyond mean performance, the variance across samples differs dramatically. SA exhibits an extremely tight distribution concentrated around *−*992. In contrast, HADOFQAOA Ideal displays a moderately wider distribution, while ibm_torino spans a broad range from *−*500 to *−*200, highlighting the impact of quantum device errors. From Figure 5 (B), it is noticeable that Unicycler formed an assembly using 509 of the 524 nodes in the string graph. However, all three solvers used only 300 nodes or less. The optimistic QUBO scores from SA and HADOFQAOA Ideal imply that the raw QUBO output from the solvers consisted of many linear nodes (one incoming and one outgoing edge) with some exception. However, from the distribution of nodes, we note that the DFS post-processing removed more than 200 nodes consistently from all the solutions. This shows that most of the sampled solutions contained long branching paths, which were simply removed from the final assembly sequence. The post-DFS node distributions further clarify qualitative differences across the solvers: ibm_torino paths are heavily truncated, rarely exceeding 100 nodes. HADOFQAOA Ideal improves on this with a peak between 100 and 150 nodes, whereas SA retains the highest structural integrity among the three, spanning between 100 and 280 nodes per sample. Figure 5 (C) is promising as we notice that even though a large proportion of nodes are left out from the final assembly, many samples still managed to form assembly sequences close to the reference Unicycler sequence. While ibm_torino and HADOFQAOA Ideal predominantly generate under-assembled contigs (peaking around 2 Mbp and 3 Mbp respectively), SA displays a distinct, high-frequency spike matching the dotted Unicycler baseline at 7.1 Mbp. However, SA also shows a long tail extending up to 9.0 Mbp, indicating that over-assembly occurs in a subset of samples due to residual cyclic paths.

**Figure 5:**
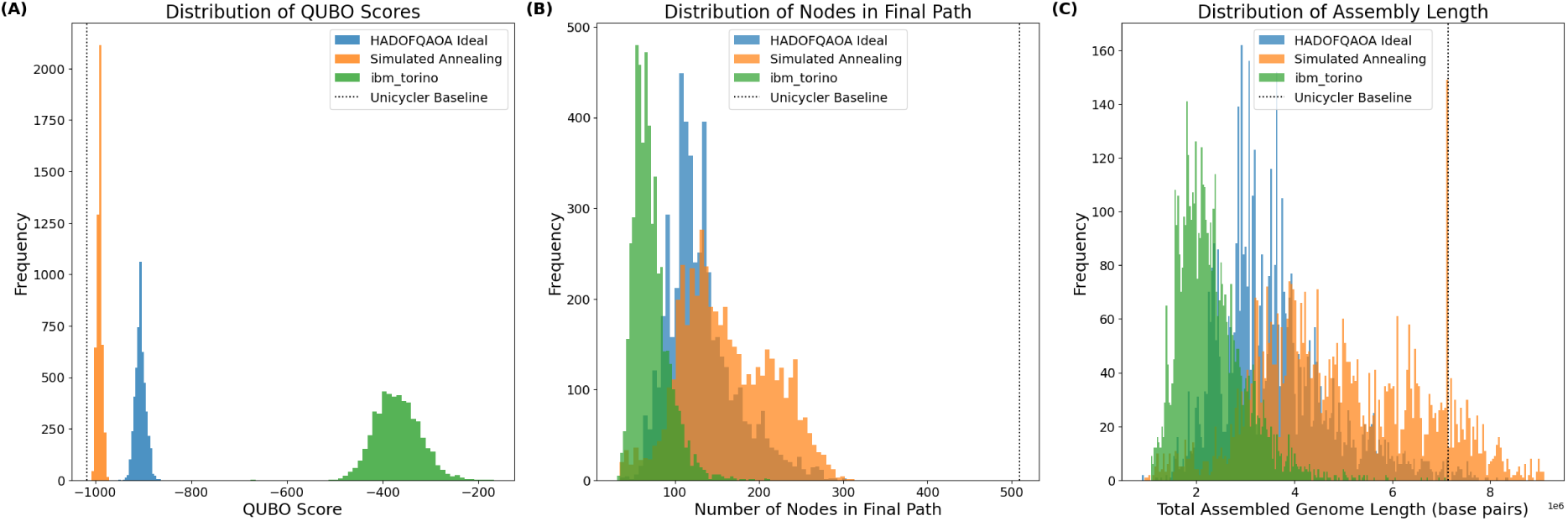
Frequency distributions of optimisation metrics from the sampled solutions of different solvers. (A) Simulated Annealing (SA) tightly converges near the Unicycler reference score (−1018), followed by HADOFQAOA Ideal, while noise on ibm_torino shifts objective values toward higher energies. (B) Distribution of retained graph nodes after Depth-First Search (DFS) post-processing. All three solvers utilize less than 300 of the total 524 nodes, highlighting significant graph pruning compared to Unicycler (509 nodes). (C) Distribution of final assembled genome lengths. SA exhibits a prominent spike aligning precisely with the Unicycler reference length at 7.1 Mbp, whereas quantum implementations skew toward shorter assemblies. Dotted vertical lines represent Unicycler reference values.

The assembly quality of the sampled solutions are evaluated using two QUAST metrics: genome fraction and duplication ratio. These metrics provide complementary measures of assembly completeness and redundancy, respectively. Taking the Unicycler assembly as the benchmarking baseline, an ideal assembly is expected to recover 100% of the reference genome while maintaining a duplication ratio of approximately 1.0. Solutions concentrated near the point (100%, 1.0) represent assemblies that provide both high genome coverage and limited redundant sequence. The distributions in Figure 6 demonstrate clear differences in assembly performance between the three solvers. The simulated annealing (SA) solutions exhibit the strongest overall distribution with respect to the two assembly metrics. A substantial concentration of samples occurs at a duplication ratio of approximately 1.0, with many of these solutions achieving high genome fractions. The distribution extends towards 100% genome fraction while remaining comparatively concentrated around the desired duplication ratio, indicating that SA consistently produced assemblies with high completeness and relatively low redundancy. The presence of a large number of solutions close to (100%, 1.0) further indicates that high-quality assemblies were not isolated outcomes within the sampled solution set.

**Figure 6:**
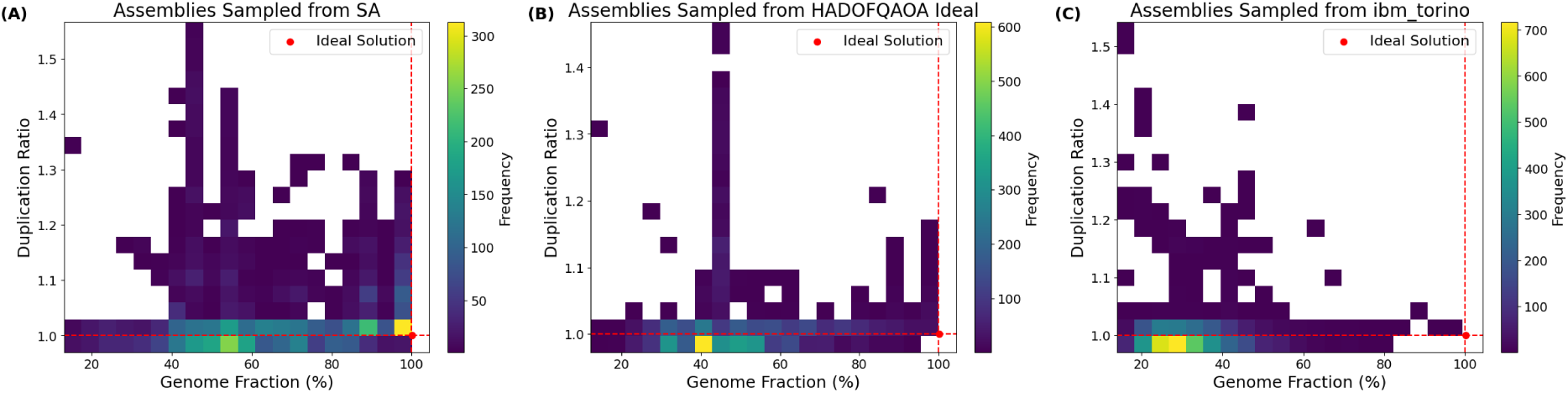
Distribution of sampled solutions from the QUBO solvers by duplication ratio and genome fraction. Since the assembly from Unicycler is considered as a baseline for benchmarking, ideal solutions must cover 100% of the genome with a duplication ratio of 1.0. The distribution of SA shows a high concentration of solutions around the ideal solution. However, for HADOFQAOA Ideal and ibm_torino the most concentrated regions cover only around 40% and 30% of the reference genome. Nevertheless, all three solvers contain at least one solution close to the ideal solution.

The HADOFQAOA Ideal distribution shows a different pattern. Although solutions reaching high genome fractions are present, the largest concentration of samples is located around approximately 40% genome fraction, predominantly at a duplication ratio close to 1.0. While the solutions in this region are not strongly redundant, they recover only a relatively small fraction of the reference genome. The distribution also contains samples extending towards higher genome fractions, including solutions close to the ideal region, demonstrating that the solver was capable of producing complete assemblies. However, these high-quality solutions represent a smaller portion of the sampled solution space than observed for SA. The real-device results from ibm_torino exhibit the most concentrated distribution at lower genome fractions, with the highest-density region occurring around approximately 30% genome fraction and a duplication ratio near 1.0. Similar to HADOFQAOA Ideal, this indicates that many sampled solutions produced assemblies with limited genome coverage despite relatively little sequence duplication. Nevertheless, the distribution extends to higher genome fractions, and at least one sampled solution approaches the ideal region. This demonstrates that the hardware execution was capable of generating highly complete assemblies, although such solutions were comparatively rare within the sampled population. The presence of solutions with duplication ratios above 1.0 with higher genome fractions also indicates that increasing completeness was sometimes accompanied by redundant sequence in the resulting assemblies. To quantify optimal solutions, we identify solutions above 95% genome fraction with a duplication ratio less than 1.05. Among the 5000 sampled assemblies from each solver, 398 SA solutions met these conditions, compared with 48 HADOFQAOA ideal simulations and only 2 executions on the real ibm_torino. Among these, 106 SA samples and 3 HADOFQAOA Ideal samples xontain 100% of the genome with a duplication ratio of 1.0, indicating perfect reconstruction of the reference genome. The highest genome fraction covered by ibm_torino samples from this run was 99.348% with a duplication ratio of 1.0. These results provide a proof-of-concept that biologically plausible genome assemblies can be sampled from the proposed QUBO formulation using both classical and quantum optimisation approaches.

### 4.4 Relationship Between QUBO Metrics and Assembly Quality for De Novo Assembly

To establish whether optimisation metrics can serve as proxies for biological assembly quality, we evaluate the joint distributions between internal optimisation outputs (QUBO score, path node count, total genome length) and reference-based QUAST metrics (genome fraction, duplication ratio) across all sample solutions (Figure 7).

**Figure 7:**
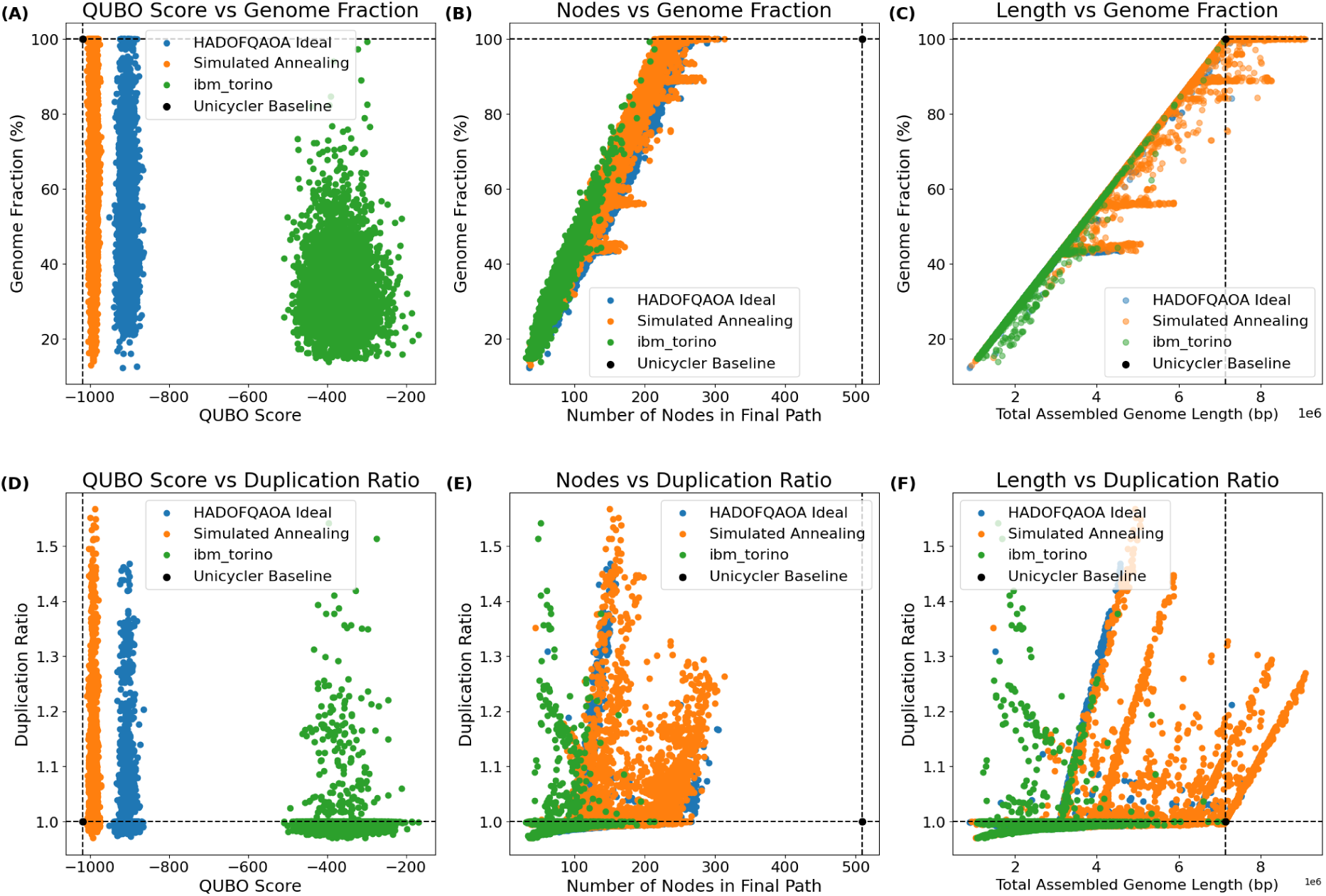
Joint distributions relating internal QUBO optimisation metrics to biological assembly metrics from QUAST across QUBO solvers. (A) QUBO score vs. genome fraction, showing a decoupling between objective minimization and sequence coverage. (B) Retained post-DFS path node count vs. genome fraction, exhibiting a strong linear correlation where trajectories with >200 nodes approach 100% completeness. (C) Total assembled genome length vs. genome fraction, demonstrating a linear increase that plateaus at 100% completeness upon reaching the Unicycler reference size (7.1 Mbp). (D) QUBO score vs. duplication ratio, where low-energy solutions display variable redundancy up to 1.6. (E) Retained node count vs. duplication ratio, with intermediate node counts showing increased variance before re-converging toward 1.0. (F) Total assembled genome length vs. duplication ratio, highlighting discrete linear trajectories corresponding to over-assembled repeat regions extending beyond the 7.1 Mbp baseline. Dashed lines and black markers denote the reference values from the Unicycler baseline assembly.

Figure 7(A) demonstrates a clear decoupling between raw QUBO score minimisation and genome fraction. Across all three solvers, solutions achieving similar QUBO scores span almost the full spectrum of genome completeness (*∼* 15% to 100%). Furthermore, lower QUBO scores do not inherently guarantee structural correctness; as shown in Figure 7(D), solutions near optimal QUBO values occasionally exhibit inflated duplication ratios up to 1.6, indicating that objective minimisation alone cannot distinguish complete single-copy traversals from assemblies carrying localised structural duplications. In contrast to the QUBO score, the number of nodes retained in the post-DFS traversal path strongly correlates with biological completeness. Across all solvers, genome fraction increases strictly linearly with retained node count (Figure 7(B)). Specifically, trajectories retaining *≥* 200 nodes consistently achieve genome fractions approaching 100%. As highlighted in Figure 7(E), intermediate node counts (100–250 nodes) exhibit higher variance in duplication ratio (1.0–1.5), whereas paths retaining *>* 250 nodes converge toward a duplication ratio between 1.0–1.3. This may suggest that focussing on increasing the node count of sampled solutions may offer higher genome fractions and duplication ratios closer to 1.0.

Total assembled genome length provides a clear structural signature of assembly completeness and over-extension. Figure 7(C) shows a direct linear relationship between total sequence length and genome fraction, which plateaus at 100% upon reaching the Unicycler reference length of 7.1 Mbp. Figure 7(F) reveals distinct, discrete linear branches radiating outward at higher genome lengths. Assemblies overextending past 7.1 Mbp maintain a 100% genome fraction but suffer from elevated duplication ratios (*>* 1.0), confirming that sequence beyond this saturation point represents redundant repeat traversals rather than novel genomic sequence. While reference-based QUAST evaluations reveal that a threshold of *≈* 7.1 Mbp and *≥* 200 nodes isolates optimal single-copy assemblies, identifying these precise quantitative boundaries remains challenging in true de novo scenarios where the exact genome length is unknown a priori.

Nevertheless, these findings highlight a path forward for reference-free sample filtering and solver optimisation. As observed in Figure 5(C), SA naturally concentrates a high density of solution samples precisely around the true target genome length (7.1 Mbp). Sampled solutions from SA also exhibit the highest frequency of samples consisting of close to 100% genome fraction and 1.0 duplication ratio from Figure 6. This population-level convergence suggests that modal peaks in path length distributions could serve as an intrinsic self-calibrating proxy for target genome size in reference-free workflows. Additionally, incorporating raw read coverage and local graph topology directly into the QUBO penalty formulation can better penalise over-traversed repeat regions, discouraging paths that extend beyond the singlecopy baseline. Enhancing HADOF-QAOA by designing assembly-specific aggregation policies may also improve quantum solution quality.

## 5 Conclusion and Future Work

This work demonstrates a quantum-assisted genome assembly workflow for a real, clinical 7.1 Mbp *of Pseudomonas aeruginosa* isolate using Oxford Nanopore long-read data within the Unicycler pipeline. By replacing the classical layout optimisation step with a QUBO formulation and deploying HADOF for decomposition, we addressed the critical challenges highlighted in Section 1:

- Qubit and Connectivity Constraints: Standard QUBO mapping makes a 524-node string graph with 2,313 overlap edges impossible to execute monolithically on NISQ processors. HADOF successfully federated this large-scale graph into execution-ready sub-problems, bypassing physical qubit and layout restrictions on the 133-qubit ibm_torino QPU.
- NISQ Hardware Noise and Circuit Depth: Gate errors and decoherence heavily degraded raw hardware outputs (ibm_torino averaged a QUBO score of *−*370 and 30% genome fraction) compared to HADOFQAOA ideal simulations (averaging *−*906 and 40% genome fraction) and Simulated Annealing (averaging *−*992). However, HADOF’s localised sub-problem decomposition reduced circuit depths, enabling ibm_torino to sample solutions reaching up to 99.35% genome fraction.
- Penalty Formulation and Soft Constraints: Rather than relying strictly on raw QUBO score minimisation, which was observed to decouple from biological completeness, we demonstrated that downstream graph traversals can recover complete contigs even when soft penalty terms produce suboptimal objective values.
- Structural Patterns for Candidate Selection: Probabilistic solvers sample a broad spectrum of traversal candidates. By analysing joint distributions between optimisation metrics and QUAST outputs, we observed key structural patterns such as post-DFS node retention boundaries and population-level length peaks - that lay the groundwork for candidate selection strategies.

Our empirical analysis revealed fundamental insights regarding optimisation behaviour and biological validity. Minimising the QUBO score alone does not guarantee a complete assembly and can occasionally favour redundant repeat traversals with duplication ratios up to 1.6. Post-DFS retained node count scales strictly linearly with genome fraction across all solvers. Retaining *≥* 200 nodes consistently yields genome fractions approaching 100%. Total sequence length increases linearly with genome fraction until saturating at the target reference length (7.1 Mbp). Assembled sequences extending beyond this point maintain 100% genome fraction but exhibit discrete linear duplication branches (*>* 1.0), signalling repeat-induced over-assembly. Beyond the results presented here, this work provides an open-source foundation for further development of quantum-assisted genome assembly. We make the HADOF implementation and the complete benchmarking pipeline publicly available through the accompanying GitHub repository^6^, enabling reproduction of the experiments and facilitating future investigations across different QUBO solvers, quantum devices, and genome assembly datasets. The supplementary material additionally contains the data collected from the experiments reported in this study, including executions performed on ibm_torino. These resources provide a reproducible basis for evaluating future improvements in quantum-assisted assembly and for investigating its transition from proof-of-concept experiments toward practical applications.

Building on our analysis, future work will focus on four primary directions. Further research is required to formalise a fully generalisable, reference-free candidate filtering framework. Validating these observed structural patterns across diverse bacterial and eukaryotic datasets will allow us to test whether prioritising higher node counts or identifying modal density peaks in sample distributions can reliably infer target genome lengths without prior reference knowledge. Reformulating the QUBO objective function to incorporate raw read sequence data and local graph topology directly into penalty weights. This is to penalise over-traversed repeat regions and suppress sequence over-extension past the single-copy baseline. Tailoring HADOF’s solution aggregation specifically for genome assembly graph structures. Currently, 5000 sample solutions are created independently from each decomposed 5-qubit circuit. They are naively aggregated to create 5000 complete solutions by merging each solution from decomposed circuits in the same sampling order. However, the distribution of the sampled sub-solutions from each decomposed circuit may contain information about optimal solutions. This could be exploited to create an aggregation policy specific to genome assembly. Incorporating explicit constraint-handling techniques within HADOF to enforce topological path validity directly during circuit execution. Constraint handling optimisation algorithms [15, 10, 12] can be tested for compatibility with HADOF.

Although current quantum hardware remains limited, the successful execution of decomposed genome assembly optimisation from a real sequencing dataset on a real IBM QPU represents a substantial step toward practical quantum bioinformatics. Most importantly, this work demonstrates that hybrid decomposition frameworks such as HADOF can bridge the gap between the theoretical promise of quantum optimisation and the practical constraints of current hardware.

## Acknowledgments

We acknowledge the use of IBM Quantum services for this work. The views expressed are those of the authors, and do not reflect the official policy or position of IBM or the IBM Quantum team. We thank Ms. Deepani Ningthoujam for assistance with the preparation and design of the figures presented in this work.

## Data and materials availability

An open-source implementation of HADOF and the complete quantum-assisted genome assembly benchmarking pipeline as a tool for future experiments and possibly practical usage. The data collected from previous runs, including experiments on a real quantum device, is made available on a supplementary GitHub: https://github.com/Namasi-PhD/Quantum-Genome-Assembly

## Footnotes

1 7.1 Million base pair genome, compared to toy problems and smaller genomes of less than 5000 base pairs commonly seen in current literature.

2 https://anonymous.4open.science/r/Quantum-Genome-Assembly-D0E9

3 https://anonymous.4open.science/r/Quantum-Genome-Assembly-D0E9

4 k-mers are a sequence of k base pairs from the sequencing reads. Here, we look for matching k-mers within two reads, which have the same sequence, ideally. Minimap accounts for sequencing errors, even if the k-mers do not completely match.

5 Coverage describes how many times a specific nucleotide in a genome is sequenced by the sequencing machine.

6 https://anonymous.4open.science/r/Quantum-Genome-Assembly-D0E9

